# Interactional roles in research meetings: a combined conversation analytic and quantitative network approach

**DOI:** 10.1101/2022.11.15.516566

**Authors:** Marloes D. A. Bet, Aniek R. Antvelink, Stéphanie M. E. van der Burgt, Saskia M. Peerdeman, Jeroen J. G. Geurts, Joyce Lamerichs, Linda Douw

## Abstract

Translational processes that facilitate effective interdisciplinary communication remain a ‘black box’. Here, we aimed to operationalize translational practices by combining conversation analysis and network science to analyze interdisciplinary research meetings. We applied conversation analysis to transcribed meetings, and identified chair, clarifying, skeptical, expert, connecting, and practical actions. For each meeting, we constructed a network (*‘meetome’*) with team members as nodes, and the number of consecutive speaking turns between members as links. We found that the relative occurrence of role-specific actions correlated with network measures. We discuss how awareness of interactional roles within meetings may help to implement a translational approach in interdisciplinary research teams.

## Introduction

As modern scientific problems are becoming increasingly complex, researchers rely more than ever on interdisciplinary teamwork to solve them. Interdisciplinarity is defined as “a cooperative team effort motivated by the need to address complex problems that cut across traditional disciplines, and the capacity of new technologies to both transform existing disciplines and to generate new ones” (CohenMiller & Pate, 2019; Lorenzetti et al., 2022). The primary aim of this approach is to achieve a solution to complex problems that cannot be tackled by one research discipline alone. It is integral to distinguish interdisciplinarity from multidisciplinarity. In multidisciplinary teams, researchers work mainly as independent specialists, with occasional overlap in roles and goals. In contrast, interdisciplinary teams collaborate on joint projects, in which boundaries between scientific disciplines are less demarcated (Chamberlain-Salaun et al., 2013; Choi & Pak, 2006). Interdisciplinarity requires not only adding the knowledge of different researchers together, but also recombining knowledge in novel ways to come up with ideas and solutions for complex issues (Vestal & Mesmer-Magnus, 2020). Further, interdisciplinarity involves negotiation between the participating fields of study, which each have their own preference with respect to definitions, methodologies and relevant outcomes (Mol & Hardon, 2020).

Interdisciplinary collaboration fundamentally relies on translational processes, which transfigure and recombine knowledge into novel approaches or ideas. Therefore, the translational approach is defined by a synthetic process in which two different ideas, that seem to be incommensurate, are integrated and together form a solution that is more than the sum of its parts (Mol & Hardon, 2020; Tuertscher et al., 2014). Translation can occur in multiple ways, but the current study will focus on translation within and between scientific disciplines. Although translation is integral for researchers to perform in the increasingly interdisciplinary climate, it remains unknown which communication patterns foster translation ‘on the ground’ to promote effective interdisciplinary teamwork. Thus, the translational approach largely remains a ‘black box’.

To some extent, translational team features have been described before. An extensive literature review identified eight key features of an effective translational team (Lotrecchiano et al., 2021). The most important facilitating factor was a strong trust in competency, benevolence and integrity of all other team members, resulting in the team being perceived as psychologically safe. Other factors included clearly demarcated team roles, team-oriented communication, shared vision, flexible and adaptive behaviors contributing to a learning environment, adequate management of meetings, and frequent interdisciplinary collaboration. However, most of these factors are quite abstract and can only be achieved at the team level, providing little guidance for individual behavior.

On an individual level, translation may be promoted by adopting a *‘translational attitude’.* Such an attitude is characterized by cognitive openness and self-awareness, passion and perseverance, and proactive facilitation of information exchange (Lotrecchiano et al., 2021). Indeed, openness, humbleness or self-awareness, and a willingness to learn and share one’s own knowledge have been shown to improve the performance of interdisciplinary research teams (Tkachenko & Ardichvili, 2020). Team members may already (subconsciously) take actions that facilitate translation, but such behaviors could also be learned or stimulated, for example through team building activities or by specific interventions (McCormack & Strekalova, 2021; Wooten et al., 2015).

While previous research has focused on which factors are necessary for translation to occur in theory, little is understood on how these key elements of translation can be established in practice. Science communication research at the level of team meetings is scarce, and therefore the fundamental processes and discourse that takes place at the root of scientific knowledge and innovation is often obscure (van der Sanden, 2016). To this end, we combine qualitative (i.e. conversation analysis) and quantitative (i.e. network science) approaches to investigate translational communication processes within research groups, with the aim to identify and describe different role patterns that may be important to achieve translation.

## Methodological framework

### Conversation analysis

Conversation Analysis (CA) is an inductive approach to analyze naturally occurring speech, that orients to all talk as actions. Therefore, CA is not focused on what people say, but on what people “do” or achieve with what they say. CA assumes that there is an underlying social organization in all talk, which allows for the investigation of interactions (Sacks, 1992). In accordance with CA methodology, the current study worked with audio data. This allowed the meetings to be analyzed in great detail as they were unfolding, as minor elements (such as pauses, or ‘uhh’s) were also captured. In this study, translation was conceptualized as actions, i.e. as something that researchers *did* in their exchange with one another. To stay close to activities as they were organized by the participants themselves, we made use of the next-turn-proof procedure. Here, the analyst checked the correctness of their interpretation against co-participants’ interpretation as expressed in the subsequent turns. These turns should provide proof that the analysts’ interpretation is correct (Sacks et al., 1974). In this context, this meant that assumptions as to whether utterances had translational properties were based on the uptake of other members of the research team. For example, when somebody used a metaphor that integrated different knowledge forms, it only was deemed to function as a translational action if others also adapted this metaphor. Consequently, translation was defined as an interactionally established phenomenon, rather than operating on an individual level.

Because of the properties described above, CA has proven itself an effective method to translate abstract communication concepts, such as for example “empathy” or “rapport” into concrete actions that happen at the conversation-level (Heritage, 2011; Prior, 2018). Therefore, CA was also determined to be an effective method to research how translation was interactionally established in research meetings.

### Network science

Most social systems are structured as a network, and can thus be elegantly modeled using network science. Network science views systems as networks, consisting of nodes that are interconnected through links (Euler, 1741), and is often used in the context of neuroscience, where the brain network is coined the ‘connectome’ (Bassett & Sporns, 2017; Kostic et al., 2020; Sporns, 2018). However, since it relies on fundamentally mathematical concepts of graph theory, network science can be applied to any other system with a network organization, of which group-level communication is a prime example (Tuertscher et al., 2014). As such, it represents a quantitative approach towards visualizing and abstracting the features of a networked system.

In the current work, research meetings are investigated according to the principles of network science, i.e. represented as a network, which will be referred to as ‘meetomes’ throughout this work; the meeting’s connectome, as it were. In such a meetome, nodes correspond to team members and links correspond to their respective interactions. In this way, network analysis provides quantitative measures of individual team members’ turn-taking during meetings as well as the general interactional organization of that meeting. Depending on the processes in the meeting, the general structure or *topology* of a meetome may vary tremendously. The topology of a network can be quantified using various properties called network measures. Network measures are mathematically defined, but do not intrinsically have an assigned meaning in the system that the networks describe. Network measures can be used to describe general meeting topology, as well as team members’ individual position in the meeting.

## Methods

### Data collection

From 2017 to 2019, 70 research meetings from the Clinical Neuroscience research team of the department of Anatomy and Neurosciences at Amsterdam UMC were recorded. This interdisciplinary research group consisted of 26 members, most of whom had a background in neuroscience. Further, team members’ expertise ranged from neuroimaging, pathology and neurophysiology, neurobiology, neuro-immunology, psychology and psychiatry, to movement and health sciences, medicine, philosophy, artificial intelligence, computational science and biomedical engineering. The research meetings of this team were held weekly, and had an approximate duration of 90 minutes.

Each of the recordings was replayed by one of the team members, who listed turn-taking by each of the team members in chronological order using time stamps. These turn-taking lists were used for quantitative network analysis. In addition, 11 of the 70 recorded meetings were manually selected and transcribed to perform qualitative CA. In selecting the 11 meetings for qualitative analysis, we aimed to pick meetings that were representative of the entire collection of meetings in terms of members present, topics covered and meeting format used. The sample was limited to 11 meetings because this number of meeting recordings sufficed for data saturation, as similar patterns were found to recur across meetings. Transcription was performed manually, because there was no freely accessible software available to automatically transcribe spoken language in Dutch with sufficient quality and resolution. For example, if a suggestion was replied to with a confirming ‘mm-hm’, then this would prove the suggestion to be considered valid by at least one team member; however, available software could not pick up on these subtle utterances. Because all meetings were in Dutch, as well as for privacy and confidentiality reasons, transcripts will not be published apart from anonymized illustrative examples in the Results section and Supplementary Materials.

For 41 meetings, an evaluative survey was distributed among all present team members right after the meeting ended. This survey included 8 items that were rated on a 5-point Likert scale (Supplementary Section 1).

### Study procedures

Transcription and exploration of the data was started simultaneously. We collectively discussed the audio material in an explorative manner, to identify actions that we believed contributed to the translational capacity of the meeting. Such actions included, but were not restricted to, directly addressing another member or the entire group with a question or task; formulating a problem or idea and inviting input; relating a proposed idea to previous ideas or own experience; asking for more explanation; proposing solutions to proposed problems; explaining phenomena, ideas or methods; being critical of someone’s statements; and many more. Using the insights from these explorations, we extracted the actions implied in speech and related them to one another to form coherent role patterns from an interactional perspective. To ensure consistency and reproducibility of our method, we implemented a systematic approach, described in the following section.

Of the 11 transcripts, 7 (~10 hours of audio material) were quantified by counting how many times each team member had produced a role-specific utterance. We constructed and employed a systematic approach to identify and quantify these actions, described in full detail in Supplementary Section 2. First, taking on a CA approach, we marked all turns that significantly contributed to a more fruitful discussion or allowed a window of opportunity for translation to occur. We identified these important utterances by using the next-turn proof procedure, as explained previously. In the context of our study, this meant that utterances were considered important if other team members reacted to the utterance as such. Thereafter, all important utterances were labeled as a particular action, e.g. summarizing, challenging the speaker, etc. Sets of actions that together formed a coherent interactional pattern were clustered into roles. We calculated the relative occurrence of utterances that corresponded to each of the identified interactional roles per meeting (total number of role-specific utterances / total number of important utterances) and per person (number of role-specific utterances / number of important utterances).

#### Inter-rater reliability

To evaluate the agreement between two independent raters (A.A. and M.B) on which utterances to mark as important, we calculated the intra-class correlation coefficient (ICC) over a dataset (one transcript) consisting of 605 utterances in total. We used a two-way mixed effects model for the inter-rater reliability, as the raters were fixed and the utterances were random. The ICC was calculated prospectively over the independent ratings of the same dataset, and retrospectively, after the raters had compared their scoring and exchanged their ideas about the meaning, function and relevance of the utterances.

#### Meetomes

We constructed a meetome for each meeting, in which nodes represented team members and links were defined by the chronological order of turn-taking in a meeting. Thus, if node A produced an utterance, and thereafter node B produced an utterance, then node A and B shared a link, irrespective of whether node B’s utterance was intended as a response to that of node A or not. Over an entire meeting, the weight of the final link between any two members represented the sum of all occurrences where node A spoke after node B or vice versa (Figure 2).

**Figure 1.**
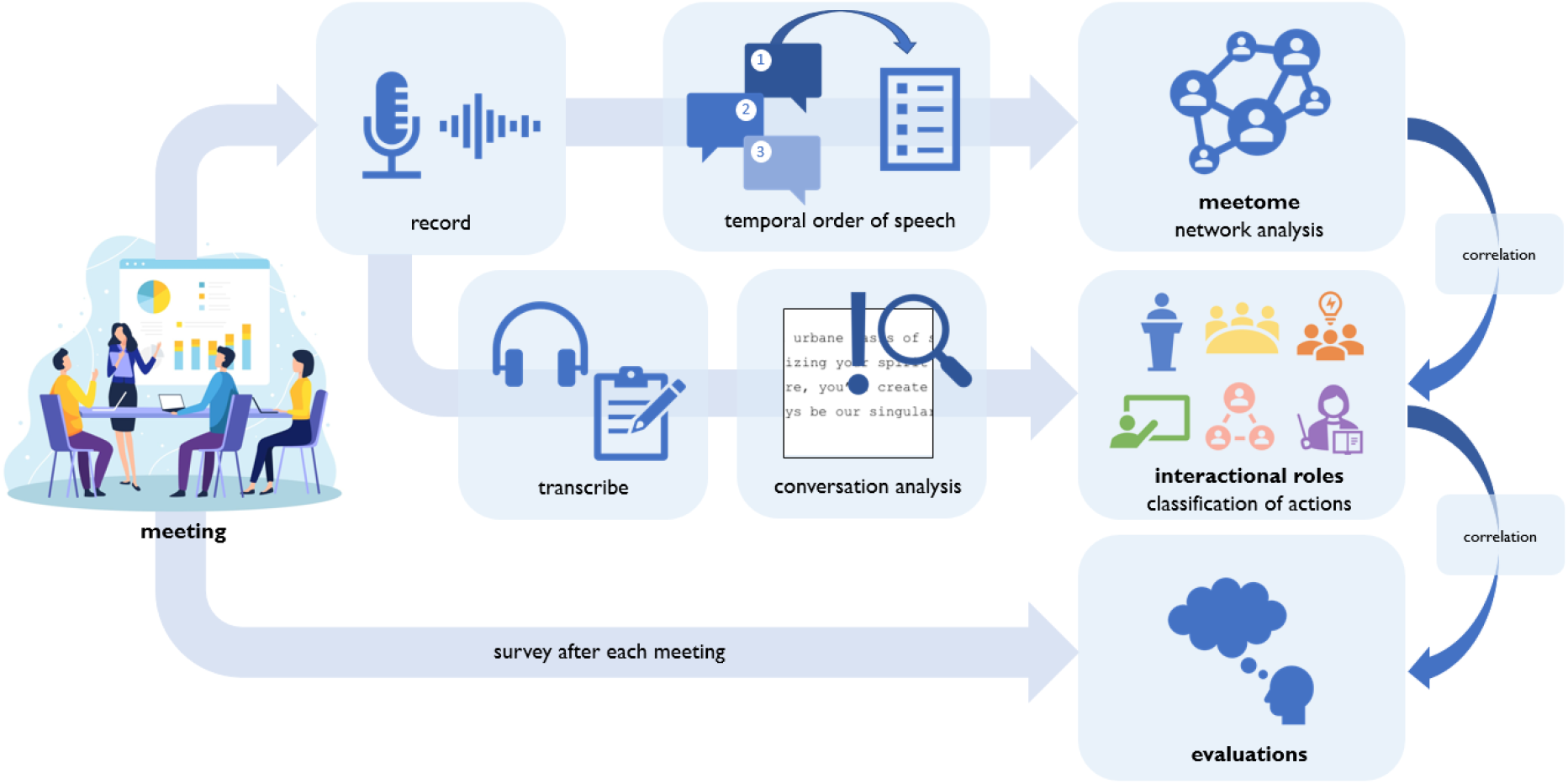
Overview of study procedures

**Figure 2.**
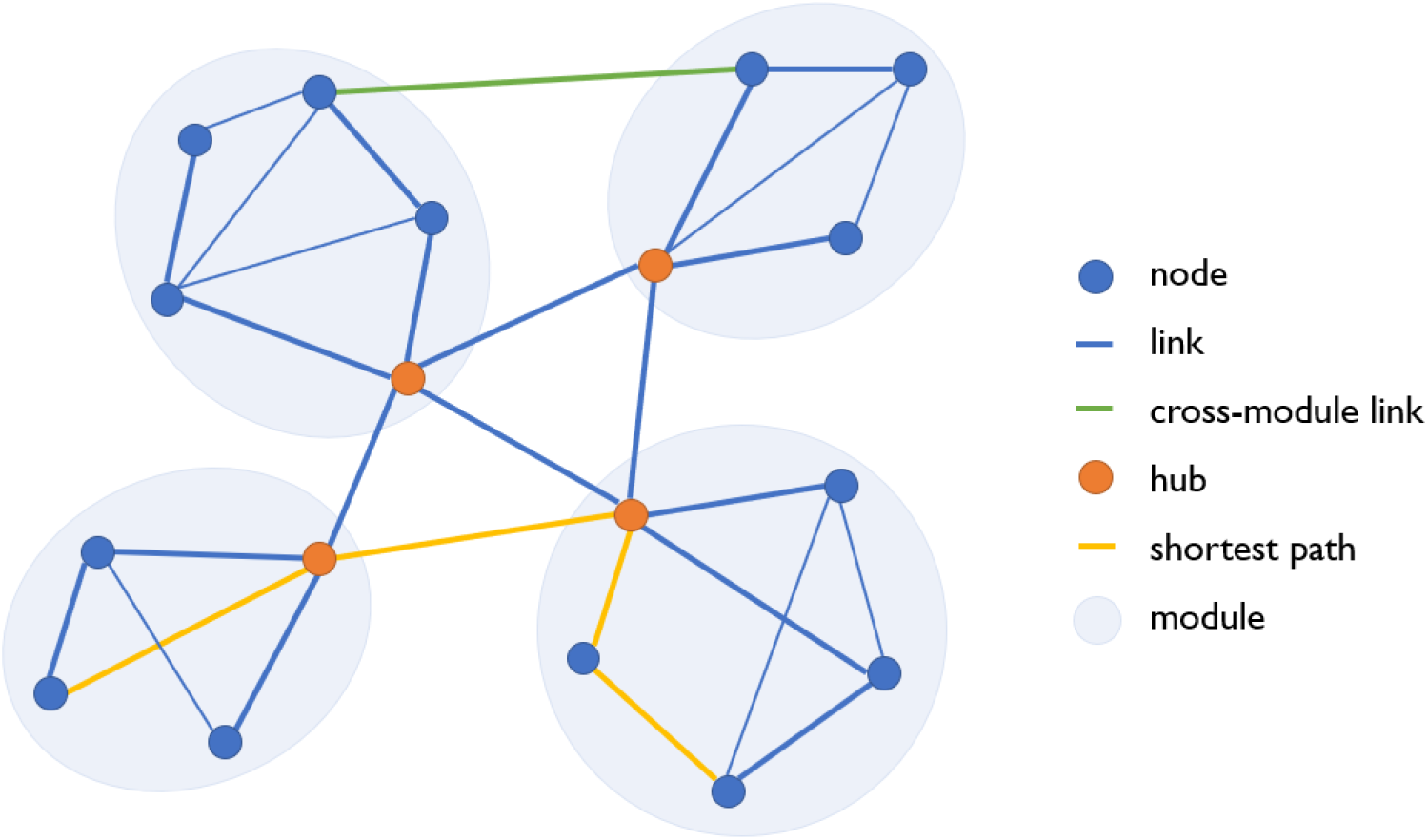
Example of a meetome. Nodes correspond to team members, and links correspond to their respective interactions, which can vary in interaction partner and in interaction strength. Interaction strength, i.e. weight, depends on the number of shared interactions and is represented here by link thickness. A node’s degree represents the number of direct connections to other nodes. **Nodal strength** is the product of degree and interaction strength, and reflects a node’s weighted connectivity. Networks tend to be organized in highly interconnected clusters of nodes, named **modules**. A node’s **within-module degree** reflects the number of links to nodes in the same module. Nodes can also link outside of their own module (*green*); the **participation coefficient** is a measure of between-module connectivity. Although the two nodes connected by the yellow path can link through other routes, the *yellow* path requires the least intermediate nodes, and thus best represents efficiency of information transfer. Efficiency can be calculated for one node in a module (**local efficiency**) or as a collective measure across the entire network (**global efficiency**). **Betweenness centrality** indicates how many shortest paths run through one node; here, the *red* nodes have a high betweenness centrality and can therefore be considered hubs that link the modules together.

#### Network measures

Throughout a meeting, not all members would follow up on each other’s turn, and therefore, some pairs of team members did not show a direct connection in the meetome. Instead, such members exchanged information via an extra step, i.e. a third member that was connected to both of them. The steps taken between two non-connected members together formed a path through the meetome (Figure 2). The length of the **shortest path** between any two members held information about the efficiency of node-to-node information flow through a network; this was most efficient when nodes shared a direct link. Therefore, we calculated **global efficiency** as the mean of all shortest path lengths taken together, representing the global capacity of a meetome to transfer information via short paths (Latora & Marchiori, 2001). In the context of science communication, this could show how much information was effectively transferred from the speaker(s) to other team members in the meeting. Whereas global efficiency measures the integration in a network, **modularity** measures its segregation. Most networks exhibit a tendency to be organized as a collection of smaller subnetworks or modules (Newman, 2006). In meetomes, modules could represent subgroups of people who have a higher tendency to follow up on each other’s turns than those of the rest of the group. This could be related to certain topics being discussed, which were relevant for some members but not for others.

In addition to global network topology, we calculated nodal network measures, i.e. local efficiency, nodal strength, participation coefficient, betweenness centrality, and within-module degree. These measures assess the network theoretical role of each team member in the meetome.

**Local efficiency** reflects a node’s integratory capacity at a local level (Latora & Marchiori, 2001). In a meetome, we interpreted local efficiency as the extent to which one person facilitates efficient information flow within their module, i.e. the group of people that they tend to interact with the most. The **participation coefficient** indicates a node’s tendency to link to nodes outside of its own module, providing a measure of between-module connectivity. Thereby, on a nodal level, it represented the diversity of team members’ interaction partners (Guimera & Nunes Amaral, 2005). Conversely, one’s importance within their own module was represented by **within-module degree**, reflecting the number of interaction partners within a team member’s own module (Guimera & Nunes Amaral, 2005). Further, **nodal strength** considers all links, and also takes into account the weights of those links, to quantify the total connectivity strength of a node in the network. A more sophisticated way to measure a node’s importance in a network is by calculating **betweenness centrality**. As explained, in efficient networks, non-connected nodes exchange information through shortest paths that run through other nodes. Betweenness centrality represents the number of shortest paths in the network that pass through a specific node, i.e. the number of other node pairs that depend on that node for information exchange (Freeman, 1978). In a meetome, a team member had high betweenness centrality if they mediated information transfer between many pairs of other people.

Any of aforementioned nodal measures can be used to determine whether a given node is a **hub**. Hubs, regardless of how they are operationalized, are key integrators of information in the brain, either by connecting different subsystems or simply by having a very high number of overall connections (Bassett & Sporns, 2017; van den Heuvel & Sporns, 2011, 2013). However, their high connectivity within the system also makes hubs the most vulnerable to disruption (Holme et al., 2002). In meetomes, we postulated that hubs were the bottlenecks of the meeting, responsible for distributing information throughout the entire network of team members. Therefore, betweenness centrality was the most suitable descriptor of hubness in this context.

### Statistical analysis

First, the occurrences of role-specific actions were counted and normalized per meeting. These action proportions per meeting were correlated to global network measures of each meeting, i.e. global efficiency and modularity, using Pearson’s correlation test.

Second, the occurrences of role-specific actions were counted and normalized per individual team member. We tested for associations between these action proportions per person and nodal network measures, i.e. local efficiency; nodal strength; participation coefficient; within-module degree; and betweenness centrality. To account for nested data (the same team members recurring in each meeting), individual-level analyses were performed using linear mixed models.

Finally, we tested for associations between normalized action proportions per person and respective normalized evaluations of each meeting. Again, a linear mixed model was used to account for nested data due to recurring team members across meetings.

For individual-level analyses, we used the function mixedlm in the package statsmodels in Python v.3.8.3. Meeting-level analyses were performed using the function corr in the package pandas. All code used for data analysis is available on Github via https://github.com/multinetlab-amsterdam/projects/tree/master/meetomes_2022.

## Results

### Conversation analytic interactional role classification

Based on CA of research meeting recordings, we identified various actions implied in speech that were recurring and in our view potentially important for translation. We grouped these actions together in six coherent communication patterns, i.e. interactional roles. Initial role patterns emerged after analyzing roughly six hours of material, and as we analyzed the remaining transcript data to expand on these patterns, no additional roles emerged anymore. All of the important utterances could be characterized as one of the actions listed in Supplementary Table 1, indicating that this provided a close to complete classification of actions and roles that were relevant for translational neuroscience, or at least in the context of these specific research meetings. The following sections provide a detailed description of each interactional role.

The most prominent role was the **chair**. The group leader typically performed most chair actions, such as taking charge of the agenda, deciding which topics were discussed, and determining who got the turn to speak. Interestingly, the person taking up these chair actions typically noticed when people got interrupted or talked over, and (re-)allocated the turn to the previous speaker, thereby establishing equivalent importance of all group members in the discussion, which appeared important to facilitate interaction between disciplines. Further chair actions were quizzing the group and delegating tasks, but also encouraging and complimenting team members on their performance, while keeping the group oriented toward long-term goals and the bigger picture. Thereby, chair actions were important determinants of the atmosphere within the research meeting, and could greatly impact perceived psychological safety within the team. Fitting with this characterization is the fact that we noticed that the chair often joked, particularly after exhibiting behavior that could be interpreted by other team members as strict, bossy or hostile.

#### Excerpt 1. Example of a chair action

> *After a long discussion that emerged following someone’s research presentation, the chair makes a move to close the subject.*
>
> Louis: “Okay Elizabeth. We have had quite some discussion within the group, and also among the department, which is very nice I think because with that you perfect your research. Do you have enough input to go on?”

In addition, two interactional roles were characterized by asking a lot of questions, but the way in which they did so was fundamentally different. On the one hand, we characterized **clarifying** actions, which were performed by the highest number of different team members. The most typical clarifying action was an open question that invited the speaker to expand on a subject, but other clarifying actions were summarizing and checking whether they understood correctly. Clarifying actions typically served to communicate that a team member had not fully understood a particular detail of what was said, and invited the speaker to elaborate. This was usually the case for members that did not share the same background as the speaker, and therefore clarifying actions were likely important for translation; before the team could bridge scientific disciplines, some common ground of knowledge needed to be reached within the entire group. Clarifying actions clearly oriented to the knowledge of the speaker, with the aim to increase the clarifier’s own understanding of the subject. In doing so, they created an opportunity for the rest of the group to ask any questions they had, or to provide their own view on the matter. This provided natural ‘checkpoints’ to summarize the information shared thus far and to ensure that the group could continue the discussion on the same level of understanding. Thereby, the borders between disciplines within the research group could begin to fade, which may have helped the team to move towards a translational approach.

#### Excerpt 2. Example of a clarifying action

> *Simone asks a clarifying question after a presentation on a new type of treatment for leukemia. Here she orients to the knowledge of the other. Furthermore, by asking this question, one of the most important aspects of this new research is highlighted, making it an effective clarifying question.*
>
> Simone: “And, do you think that, how, kind of, how much more effective [than chemo-treatment] would this [treatment] be, you think, or would you not know?”

On the other hand, we noticed **skeptical** actions, often characterized by challenging or even confronting questions, and a skeptical attitude. These questions were not meant to further a team members’ own understanding of the subject matter, but rather served to test the speaker on their knowledge of the subject, the validity of their research methods, and the implications of the results. Skeptical actions were often performed by more senior members of the group, and served to test research quality against their typically sizable knowledge base. The speaker would typically react by going in-depth on the details, and thereby skeptical actions regularly sparked a vivid discussion within the entire group. Such discussions could be beneficial for translation, and additionally provided an incentive to the entire group to think critically about the research, thereby ensuring high quality of research output. Yet, skeptic actions could challenge the credibility of the speaker and occasionally catch them off guard, thereby posing a potential threat to perceived psychological safety.

#### Excerpt 3. Example of a skeptical action

> *After a colleague has presented their research, the skeptic intervenes by saying that some of the terms used can be interpreted in many ways and asks for a specific definition.*
>
> Steven: “You use a lot of container concepts, right. Lesion is a typical container concept. Of course, a lot is packed in there, but the question remains, what do you mean with lesion?”

Fourthly, we observed instances of **expert** actions. Whenever a speaker struggled to answer a question, or when there was general confusion on the state of affairs during a meeting, typically a senior team member would intervene to explain the topic and answer the question under discussion. These expert actions served to dissolve the confusion around the subject and thereby the accessory commotion. Notably, the expert was usually not allocated a turn by anyone, but acted on their own accord. Other expert actions included correcting others or even criticizing their statements, but this was done in a very matter-of-fact way, clearly distinct from skeptical actions. Namely, expert actions served not to test other team members but to clarify a subject for the entire group. If successful, the rest of the group would acknowledge the expert’s superior knowledge and accept their explanation as the definitive answer to the question at hand. However, we also observed utterances that appeared to be aimed at resolving confusion, but were apparently unsuccessful, because they sparked a discussion in the group and thereby created even more commotion. Conversely, successful expert actions could close conversation topics and allowed space for new topics to emerge in the meeting, all while teaching the group more about the subject under discussion.

#### Excerpt 4. Example of an expert action

> *When Stacy is explaining how she can calculate Z-scores for her research, Lorie, in an expert-role, steps in to slightly correct Stacy to prevent confusion for the group. Later, she even asks Stacy to explain what the pros and cons are of the two methods, to ensure that these are also clear for the rest of the group.*
>
> Stacy: So, the local method uses regressions, and the clinical method is, well, is it’s its whole own thing.
>
> Lorie: Well, it’s just, they both generally use regressions, but here we often use local control measures. […]
>
> Stacy: Yeah, so solely to just make a choice [regarding] which method we will use to do this,
>
> I got the correlation coefficient-
>
> Lorie: And could you maybe still explain the pros and cons of each method? Because I am still not sure if everybody has a clear picture of the difference.

We have also observed a few actions that we linked to a **connecting** role. This role’s main aim is to connect, in the broadest sense of the word. Connecting actions included involving others into the conversation, by asking directed questions, inviting them to provide their input, or even by explicitly allocating turns. But connecting actions could also concern information; for example, coming up with metaphors or other innovative ways of understanding, or pointing out parallels between different subjects. In the context of research, connecting actions could emphasize similarities and dissimilarities in methods and conceptual frameworks of studies, and relate new findings to previous literature. Connecting actions associated striking aspects of a story to previous knowledge and sparked broader thinking and creativity in the group. This ability to facilitate exchange between known and new information may contribute to information exchange between scientific disciplines, and drive interdisciplinary teams to work in a more translational way. We even observed one instance where someone proposed to introduce another team member to an acquaintance of theirs, so that the two could collaborate on a topic that they both were working on. This showcases that connecting actions are integral to build the bridges required for effective interdisciplinary teamwork. However, connector actions were least frequently observed in our analyzed meetings.

#### Excerpt 5. Example of a connector action

> *Will is discussing some of the results of his research. Here the connector, Leonard, jumps in, to propose a possible conclusion of some of his results. Will’s answer showcases that Leonard remark has provided him with new insights into his research.*
>
> Leonard: So, since that isn’t the case, you could say that the immunization doesn’t have an effect on the status of your synapses.
>
> Will: Ehm… Yeah, yeah, I’d never thought of it like ehh… Yeah, never really thought about it, but yeah, maybe yes.

Finally, we have observed some instances of **practical** actions. Such actions were very pragmatic and solution-oriented, and had some similarities to expert actions; both require the team member to be knowledgeable on the subject, and both types of actions are intended to help the speaker. However, whereas expert actions helped to clarify a topic or a problem, practical actions were characterized by proposing a concrete solution to the issue at hand. Such actions usually occurred when the speaker proposed a problem, expressed a concern or explicitly asked the group for feedback on how to continue. Practical actions could take the form of advice (‘I would…’), tools (‘I have a script that does that…’), an analysis of pros and cons of different solutions (‘You could do it like this, but then you would compromise on accuracy…’), or proposing step-by-step solutions (‘If you do X, you can then do Y and thereby achieve Z’). In contrast to expert actions, suggestions arising from practical actions need not be accepted as the go-to answer by the rest of the group, but were solely oriented to the team member that introduced the problem. An important but implicit function of practical actions was to reassure the speaker that their problem could be fixed, and that they would be helped. So besides making research practices more effective, practical actions could contribute to a sense of trust and solidarity in the group, increasing psychological safety, which is an important prerequisite for translation.

#### Excerpt 6. Example of a practical action

> *After someone has suggested Elizabeth could use a specific technique to map her results, she signifies that she does not fully understand how to do that yet. Here Will, performing a practical action, steps in to offer help. With Elizabeth’s acceptance the topic is closed, and, hence, it does not need to take up valuable group discussion time anymore.*
>
> Elizabeth: What do you do then?
>
> Will: We can have a look at it at it this afternoon.
>
> Elizabeth: Okay, nice.

Just as actions were not restricted to any single person, team members were not bound to one role pattern, but were dynamic in the types of actions that they performed (Figure 3). However, we did notice that most team members showed a preference for a specific role. Moreover, roles and corresponding actions also depended on the topic under discussion and the patterns of hierarchy and group dynamics. For instance, when there was a pressed or tense atmosphere, often only senior members would speak. Then, after someone made a joke, the atmosphere was lightened and more junior members would join in again. However, this interaction pattern was not explicit and distinct enough to be considered a separate role in our classification.

**Figure 3.**
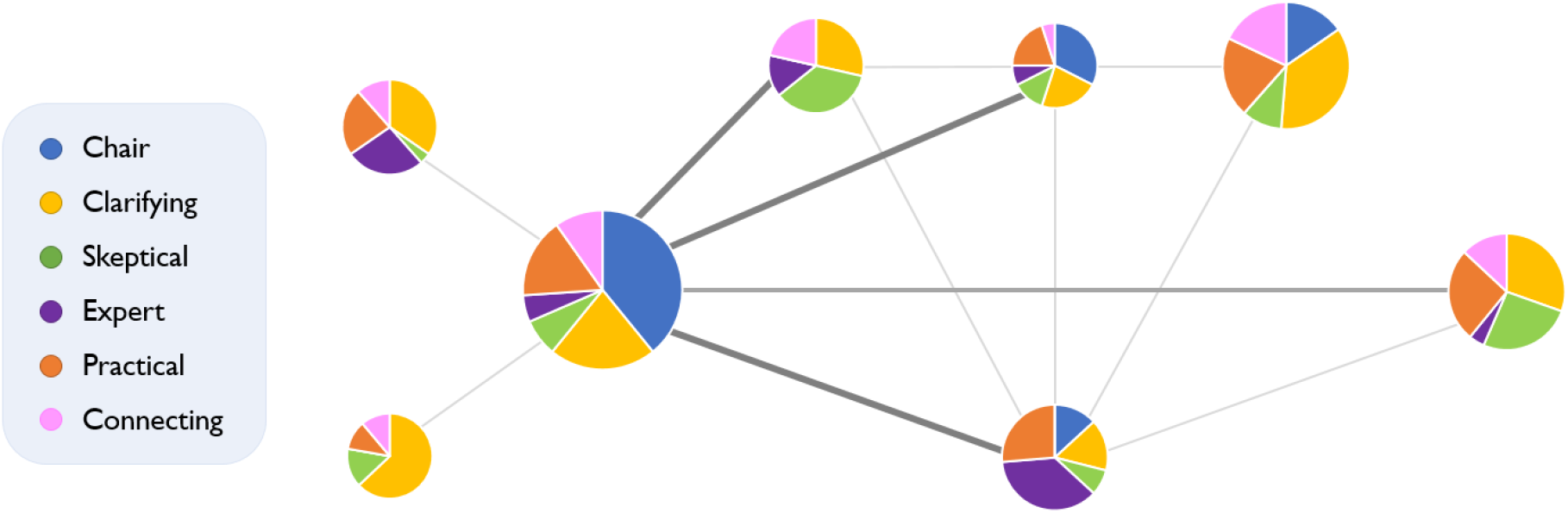
Meetome including interactional roles. Here, the eight most prominent team members from our data are visualized in an average meetome over all 70 recorded meetings. The pie charts represent the relative occurrence of role-specific actions per member across the six transcribed meetings, as determined through conversation analysis. Node size corresponds to the average nodal strength of each team member across all 70 meetings. Thickness of the links corresponds to the frequency of interaction between two nodes across all 70 meetings; thicker lines signify higher average interaction strength.

Of note, the actions described above are not always conducive for translation. All “work” being done in conversations is interactive, and which actions end up having the desired effects is determined by the subsequent terms of the other team members. For instance, not all critical questions ensure high quality research output. For example, when they are too detail-oriented, they can slow down the conversation and consume precious meeting time over minor issues. Hence, while we found that the interactional role of the skeptic was imperative in meetings, not every critical question that they posed was typified as a skeptical action. The actions belonging to the interactional roles should, thus, not be envisioned as a fitting interpretation for every turn in a conversation. Rather, they reflect an interpretation for the turns in a conversation to which the other members respond in a way that fosters translation.

#### Inter-rater agreement

Our conversation analytic approach initially yielded an ICC of 0.765 (95% CI 0.736-0.791). Thereafter, the authors (A.A. and M.B.) discussed with each other their motivations for marking utterances as important, and if they agreed, adjusted their classifications accordingly. Thereafter, the ICC increased to 0.966 (95% CI = 0.962-0.971). Strikingly, before conferring with one another, there was already a 100% consensus on which of the actions marked as important belonged to which interactional role pattern.

### Network analysis

After having quantified the relative proportion of actions corresponding to each of the identified role patterns, we correlated this to both global (global efficiency and modularity) and nodal network measures (local efficiency, nodal strength, participation coefficient, within-module degree, and betweenness centrality). At the global level (*N* = 7), strong chair-like presence, i.e. high relative proportion of chair utterances, was associated with lower global efficiency (*r* = −.683, *p* = .006) and with higher modularity in the meetome (*r* = .814, *p* = .005). Similar patterns were observed for expert actions (global efficiency: *r* = −.501, *p* = .046; modularity: *r* = .717, *p* = .020) and practical actions (global efficiency: *r* = −.575; *p* = .033; modularity: *r* = .717, *p* = .013). However, after Bonferroni correction for multiple testing, none of these correlations remained significant. Supplementary Table 2 contains the statistical parameters for all Pearson correlation tests.

At the nodal level (*N* = 89), the number of chair actions was associated with higher nodal strength (*β* = 1.211, *p* = .012), higher within-module degree (*β* = 1.525, *p =* .006), and higher betweenness centrality (*β* = 3.328, *p* < .001). Skeptical actions were associated with higher local efficiency of the corresponding team member’s node (*β* = .805; *p* = .048). Finally, connecting actions were associated with higher local efficiency (*β* = .1.288, *p* = .012), as well as with higher nodal strength (*β* = .784, *p* = .025). These associations are presented in Figure 4. After Bonferroni correction for multiple comparisons, however, only the correlation between chair actions and betweenness centrality remained (*p* < .001). Supplementary Table 3 contains the statistical parameters of all linear mixed models.

**Figure 4.**
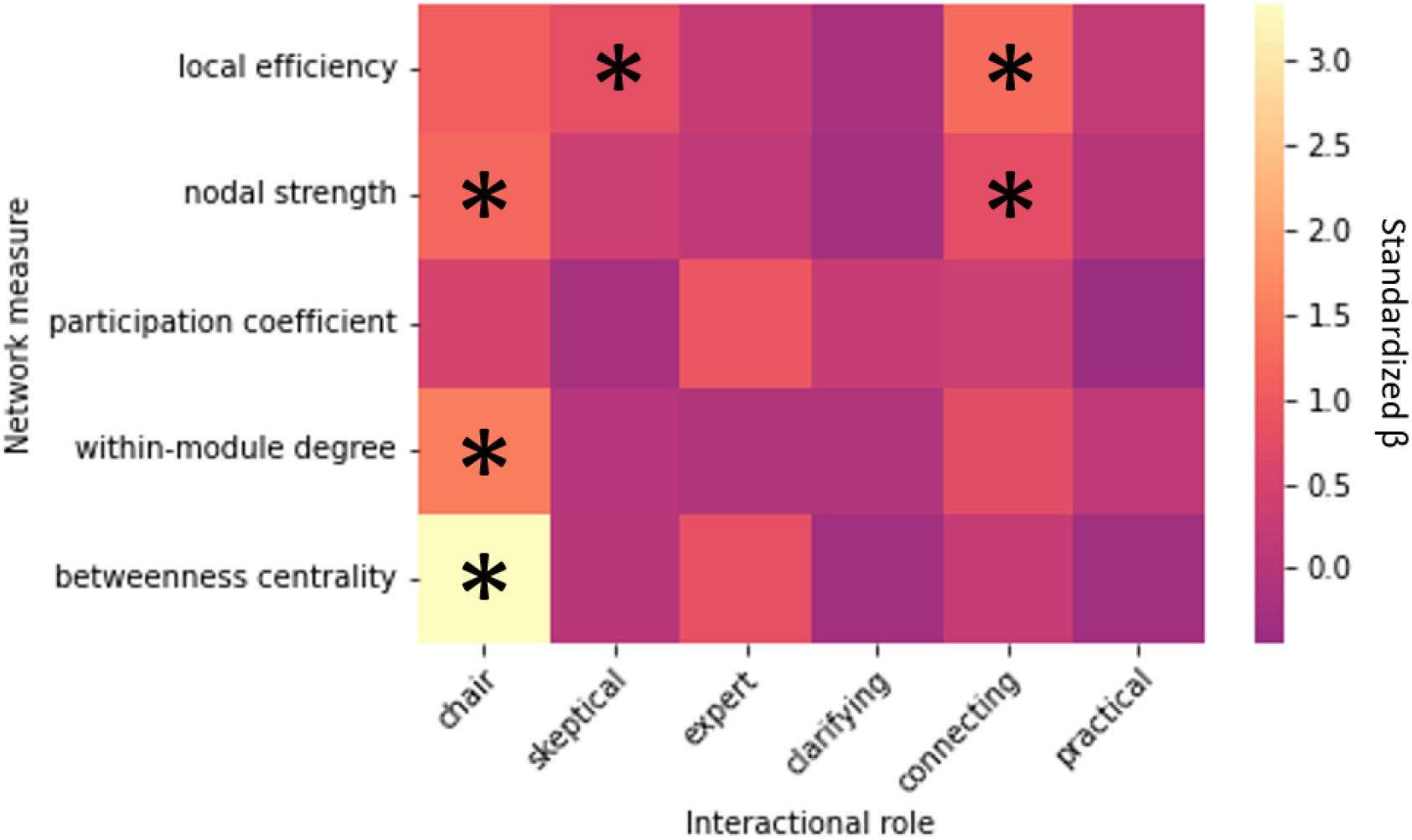
Associations between nodal network measures and relative proportions of actions.

### Correlations with meeting evaluations

To evaluate the impact of occupying each of the roles on the perception of the meeting, we correlated team members’ meeting evaluations to how often they relatively exerted a role-specific action during the meeting (*N* = 55). We found that people who exert more chair actions tended to rate the overall meeting experience more positively (*β* = .551, *p* = .036), and that people who exert more practical actions tended to rate the extent to which the meeting sparked a feeling of connection with the group more negatively (*β* = −.305, *p* = .014). However, these correlations did not persist after Bonferroni correction for multiple comparisons. Supplementary Table 3 contains the statistical parameters of all linear mixed models.

## Discussion

The aim of the current work was to identify coherent patterns of communication and thereby explore factors that may contribute to more effective translation in interdisciplinary research teams. We defined six interactive role patterns based on conversation analysis of team meetings, thereby providing a conversational typology of communicative behavior. These roles, characterized by chair, clarifying, skeptical, expert, connecting, and practical actions, provide a framework of behaviors that may contribute to translation at the root of scientific collaboration, namely in small research teams. Further, we demonstrated that the identified actions related to the network topology of the meeting, i.e. the meetome.

By describing interactional roles, we showcase the potential of specific actions to support translation in research meetings. For example, chair actions and practical actions can contribute to a psychologically safe environment, while clarifying and expert actions can establish a common ground of knowledge, skeptic actions can improve quality of research output, and connecting actions can help to bridge different knowledge bases. Because each of the interactional roles may exert a multitude of effects on the meeting, they could greatly benefit translation if used in harmonious synergy. Although certain behaviors may be determined by factors such as personality, seniority, and group dynamics, we also present behaviors that can be taught, which may help teams to adopt a translational approach. Awareness of one’s actions and their potential effects on the meeting may therefore help interdisciplinary research teams to move towards a translational approach.

Although previous research on the translational approach within research teams is scarce, various studies have identified factors that allow for effective interdisciplinary teamwork. These studies converge on the fact that team composition should be balanced in terms of scientific background (Nancarrow et al., 2013; Tkachenko & Ardichvili, 2020). If team members’ background is too similar, their marginal knowledge is largely redundant, but if they share very little expertise, then they will not possess the shared knowledge necessary to absorb and make use of one another’s expertise (Vestal & Mesmer-Magnus, 2020). Thus, in teams with diverse backgrounds, clarifying and expert actions may be crucial to establish a common ground of knowledge. This is in line with our observations that clarifying and expert actions served to explain topics that were new or unclear for a subset of the team members. Further, effective teams need a supportive, encouraging and facilitating leader who listens to team members and establishes a clear vision for the team (Lorenzetti et al., 2022; Salas et al., 2005; Tkachenko & Ardichvili, 2020).

One of the most important prerequisites for translation is a psychologically safe environment with mutual trust and solidarity (Lorenzetti et al., 2022; Salas et al., 2005; Tkachenko & Ardichvili, 2020). Such an environment can be created through some of the actions described here. For example, clarifying actions increase autonomy and credibility of the speaker, and simultaneously create an opportunity for other team members that do not fully understand everything to ask their questions. Further, practical actions show solidarity in the team and a willingness to help each other out. Related to this is the importance of collective trust in all team members’ benevolence and competency, which could be greatly promoted through expert and skeptical actions.

The correlations with network measures indicate that the actions we identified were reflected in the meetomes, and thus, that the qualitative content of speech reflected the general turn-taking structure of a meeting. For instance, chair actions were correlated with higher nodal strength, indicating many strong links to various team members; this makes sense, as most chair actions are focused on allocating turns and ‘managing’ the meeting, which requires frequent involvement in the conversation. The same is true for connecting actions, which link different topics or even team members together, and therefore also require high involvement in the meeting. Further, both skeptical and connecting actions correlated with higher local efficiency, indicating that these interactional roles facilitated efficient information transfer within their module. Chair actions were correlated with within-module degree, indicating that team members who exerted many chair actions, typically related to leadership, interacted with a higher number of people within their module. Finally, chair actions showed a particularly strong correlation with betweenness centrality, suggesting that exerting such actions may increase one’s likelihood to be a hub in the meetome. This may indicate that the chair facilitates a lot of information transfer within the meeting. However, no causality can be inferred from these associations.

Interestingly, multiple interactional roles showed the same pattern with regard to global network measures, tentatively indicating that key actions in the meeting, which could be important for translation, may be facilitated by a specific ‘translational’ meeting structure. However, global-level correlations did not survive Bonferroni correction for multiple testing, so these findings require replication before any such conclusions can be drawn. Although clarifying actions did not show significant correlations to global nor nodal network measures, we still consider such actions to be important for translation, as we observed that they allow for psychological safety while simultaneously creating an opportunity for the entire group to get on the same level of understanding.

We constructed a systematic approach to apply conversation analysis to research meetings in such a way that we could identify behaviors important to translation. Considering the extensiveness of the dataset, the agreement between the two raters using this method was remarkably high. This demonstrates that using our method, important actions could reproducibly be identified and related to a transcending interactional role patterns. This method may be used in future research to further investigate translational communication within research meetings.

One important connotation to our findings is that interactional roles are not necessarily restricted to one team member. Although we noticed that most team members showed a preference for specific actions, people were dynamic in the roles they took on, and multiple team members could even work together to perform a certain action. Which types of actions a team member performs may depend on their background, seniority, affinity with the topic under discussion or their relationship to the speaker (e.g. supervisor), among other things. Thus, in future analyses, it would be interesting to take such factors into account, for example by creating meetomes per topic rather than over the entire meeting, in which multiple topics were discussed. Not only do team members show a preference for one specific role, there are also clear individual differences between the ways in which team members fulfill a certain role. For instance, one team member typically preferred the clarifier role, but acted as an expert on occasions where more junior members were interrogated. They would take on the expert role to defend and support the junior members. Such an instance occurred twice, and we did not see any other team member use the expert role in this way. Therefore, it is important to keep in mind the individual differences that add on to this crude classification of people’s actions and contributions to the meeting.

Further, some actions were not sufficiently pronounced to be captured in an interactional role. For example, in most meetings, an appointed team member would present about their work, and this may have influenced how other team members behaved. However, as we found that the presenter was not consistently characterized by specific types of actions, we did not describe ‘presenter’ actions in our results, nor did we account for such actions in our analyses. Further, an ‘organization joker’ has previously been described as an assumed social position that propagates humor within the workplace. This has been described to relieve tension and to (temporarily) subvert hierarchical patterns, among other things (Plester, 2015; Plester & Orams, 2008). In our meetings, we noticed that particularly people performing chair actions occasionally used humor to lighten the atmosphere from time to time. However, because humor was used by many team members for many apparent purposes, and was very context-dependent, we did not characterize the ‘joker’ as an independent interactional role. Still, the role of the joker may be important for patterns of communication within research meetings, and warrants further investigation.

Since we did not evaluate any performance-related outcomes, and there was no significant correlation between any interactional role and the extent to which team members perceived the meetings to be translational, we could not demonstrate a direct and explicit link to translation here. Still, the actions we identified could indirectly contribute to translation in a context-dependent manner. Other limitations to the current work were inherent to our data. We used meeting recordings from only one research team, which limits the generalizability of our results to other teams that may operate in different ways. For example, in this research team, it was not customary for students doing an internship in the department to be present in the meetings; this may very well impact the behavioral patterns in these meetings. Moreover, since we adopted a manual approach to transcribe and analyze the data, the volume of our CA data is limited (~10 hours). This is because compared to automated transcription software, manual transcription of the recordings yielded much more accurate transcripts, in which the uptake of actions by the rest of the group could more effectively be evaluated. After having analyzed six meetings, no new interactional roles emerged, indicating that this data volume was already sufficient to recognize distinct and robust communication patterns. Still, this limited data volume does constrain the power of the meeting-level correlations of roles to global network measures. Finally, the meeting evaluations obtained from team members were very heterogeneous; for every meeting, ratings on each survey item ranged from the minimum to the maximum score. This may explain why we did not find any significant correlations between the identified interactional roles and meeting evaluations.

Therefore, in future studies, larger volumes of data should be analyzed, possibly using automated transcription software to increase efficiency. Further, it would be interesting to compare various research groups in terms of the interactional roles that emerge, and to link the relative occurrence of role-specific actions to the performance of interdisciplinary teams, particularly concerning translation. The operationalization of translation remains a prominent issue. By uncovering the communication mechanisms that may support translation within research meetings, we may begin to establish evidence-based research practices to optimally support interdisciplinary research teams.

## Conclusion

By uncovering the foundations of translation in communication, research teams can more readily take action to adapt to the increasingly interdisciplinary research climate. Here, we identified actions implied in speech that may support translation, and clustered them in interactional roles. We showed that the occurrence of such actions in the meeting are associated with the general structure of that meeting. This may provide insights into how a meeting should be structured to optimally facilitate translation. Moreover, we described interactional roles in terms of concrete actions that can be taught, and therefore may prove important for the implementation of a translational approach within interdisciplinary research teams.

## Supporting information

Supplementary Section 1

Supplementary Section 2

## Author Contributions

Authors M.B. and A.A. contributed equally to this work. All other authors have participated in the conception of the study, the data analysis, and editing of the manuscript.

## Acknowledgements

We thank all members of the Clinical Neuroscience team between 2017-2019 for participating in this study, namely Anand J. C. Eijlers, Antonio Luchicchi, Benthe Westerik, Bert A. ‘t Hart, Bram Platel, Geert J. Schenk, Gerrit Glas, Hanneke E. Hulst, Jolanda Derks, Kim A. Meijer, Laura E. Jonkman, Lieke L. Hoyng, Marijn Huiskamp, Marike van Lingen, Martijn D. Steenwijk, Matthijs J. Keijzer, Menno M. Schoonheim, Myrte Strik, Piet M. Bouman, Quinten van Geest, Shanna D. Kulik, Stefanos E. Prouskas, Svenja Kiljan, Tianne Numan, Léon C. de Bruin, and Tommy A. A. Broeders.

