## Supplementary Section 1 for "Interactional roles in research meetings: a combined conversation analytic and quantitative network approach"

**Supplementary Section 1. Survey to evaluate research meetings**

1. How satisfied are you with the meeting's overall value in helping you improve your on-the-job effectiveness?
2. How satisfied are you that the meeting was well worth the investment?
3. How satisfied are you with the overall meeting experience?
4. How satisfied are you that the meeting was motivating to you personally?
5. How satisfied are you with the quality of the educational value of the meeting?
6. How translational do you think the meeting was?
7. To what extent did the meeting spark a feeling of connection with the rest of the group?
8. To what extent were you able to understand the topics that were discussed in the meeting?
