## Supplementary Section 2 for "Interactional roles in research meetings: a combined conversation analytic and quantitative network approach"

**Supplementary Section 2. Demonstration of conversation analytic approach**

This fragment runs from 1:37:48 till 1:38:20 (h:min:sec). The meeting ends at 1:40:22.

The analysis below showcases why we have interpreted Simone's questions in this instance as a prototypical action of a **clarifying role**. We came to this decision by analyzing the interaction in great detail, according to the conversation analytic method.

Before the start of the excerpt below, Leonard has just presented his research plan on a new type of treatment against acute leukemia.

```
1   Simone:    En hoe verwacht je dat
               And how do you expect that
2              >want wat is<
               >cause what is<
3              weet je wat is nu de behandeling van eh: leuke: acute: leukemie
               do you know what is the treatment now against eh: leuke: acute
               leukemia
4              is er überhaupt iets wat ze kunnen doen wat (.) sneller werkt?
               is there even something that they could do what (.) would work
               faster?
5   Leonard:                                                    [ja eh::
                                                                [yes eh::
6   Simone:    gewone chemo?
               normal chemo?s
7   Leonard:          [volgens mij ook eh:: (.) chemo eh::
                      [i think also eh:: (.) chemo eh::
8   Louis:                                  [chemo
                                            [chemo
9   Leonard:   ja
               yeah
10  Simone:    en denk je dat
               and do you think that
11             hoe (.) zeg maar hoeveel effectiever zou dit zijn
               how (.) like how much more effective would this be
12             denk je?
               you think?
13             of heb je echt geen idee
               or do you really have no idea
14  Leonard:   nee, ik heb eh:
               no, i've got eh:
15             kijk (.) die ehh die aanpak is eh niet getarget
               look (.) that ehh that treatment is eh not targeted
16  Simone:                                                [hmm
                                                           [hmm
17  Leonard:   en dit wel
               and this is
18  Simone:          [ja:
                     [yeah:
19  Leonard:   dus ik denk zomaar dat en de doses nog lager kan
               so I think just that and the doses can be lower
20  Simone:    hmhmm
               hmhmm
21  Leonard:   en dat de effectiviteit veel hoger is
               and that the effectiveness is much higher
22             kijk eh ik eh:: ik geloof hier wel in hoor
```

***look* eh i eh:: i do believe in this**
23 eh::
***eh::***
24 Simone: hmm
***hmm***
25 James: [systemische belasting is gewoon veel lager
***[systemic burdening/taxation is just much lower***
26 Simone: hmm
***hmm***
27 Leonard: ja precies
28 ***yes exactly***

Simone starts her inquiry in line 1 with ‘and how do you expect that’, a sentence she then breaks off. This probably refers to the eventual line of her questioning, namely how Leonard expects this treatment to compare to other leukemia treatments in terms of effectiveness (line 11-13). Consequently, she then starts her inquiry into what the current treatment is. Here she orients to Leonard’s knowledge by using the words ‘do you know’ (line 3), characteristic of a clarifying action.  While Leonard does not adapt a prototypical hub in this instance, he plays an important role in this conversation as presents his own research, the topic currently at hand. Therefore, it is logical that he is treated as the authority on the subject. In line 4 Simone expands her turn by asking the follow-up question whether there “even” is a possible faster treatment. Through her phrasing “is there even something” (line 4), the question favors a negative response.  When Leonard starts to answer (line 5), Simone interrupts him to provide an answer herself (line 6). In doing so, she is showcasing that she has some knowledge on the subject, while at the same time leaving Leonard to confirm her answer and consequently treating him as the authority on the subject. Leonard, however, responds somewhat hesitantly using the words “i think”, producing multiple “eh::”’s and multiple small pauses.

At this point Louis, the chair of the meeting, steps in to confirm both Simone’s and Leonard’s answer. This is interesting as it also exemplifies the position the chair holds in a meeting. Here it can be seen that the chair feels comfortable stepping into a conversation in an expert position, intervening in a conversation between Simone and Leonard to answer a question posed to Leonard. That his answer is accepted by both parties can be seen by Leonard confirming his answer (line 9) and by Simone continuing her line of questioning (line 10-13) based on this now established fact.

When asking her follow-up questions Simone again orients to Leonard’s knowledge by using “[do] you think” twice. Furthermore, by producing the tag question in line 13 she phrases her question non-optimally, providing more room for Leonard to provide a negative answer. Heritage (2004) describes that in hospital settings some questions are phrased optimally, e.g.

“Are you married?”, whereas others are phrased non-optimally, e.g. “Are you divorced?”. The first question allows the person to confirm good news by responding affirmatively, whereas in the second instance the person has to reject the supposition inherent in the question to confirm good news. The question in line 13 is also phrased non-optimally as an affirmative response confirms the ‘bad’ news that Leonard “really ha[s] no idea” how effective the new treatment is. This phrasing then makes it easier for Leonard to admit a lack of knowledge on that aspect of the treatment.

It is also interesting to note, however, that by placing emphasis on “how” twice in line 11, she seems to be inquiring about specific data about how much more effective this treatment is, rather than a general statement. This specific inquiry is also reflected in Leonard’s response. It can be seen that he starts his response with “no I’ve got eh:, look (.)” (line 14-15), a preface such as “look” usually signals that a negative response is coming. While his answer is not negative in that sense, as he does indicate that he knows that this treatment is more effective, he cannot provide specific data on the effectivity. Consequently, he proceeds to explain the principle that allows this technique to be more effective. Throughout his answer, Simone produces multiple continuers (line 16, 18, 20 and 24). By doing so, she is signals active listening on her part (Fitzgerald & Leudar, 2010).

Eventually, considering the lack of specific data, Leonard appeals to his reputation as researcher and colleague in arguing that this new treatment is more effective. By saying “look eh i eh:: i do believe in this”, he treats his own believing as something that carries epistemic weight. At this point James steps in with a brief summarizing action (line 25). He does not provide new information but instead summarizes Leonard’s account in more succinct terms, and thereby also strengthens Leonard’s account. Space does not allow us to show the subsequent turns of this conversation, however, here in it becomes apparent that James does so to wrap up the conversation between Leonard and Simone. Subsequently, he uses the opportunity to introduce his own question immediately after this extract ends. Leonard confirms James’ summary with “precisely” in line 26. This word not only confirms James’ account, but also upgrades Leonard’s epistemic position, signaling that Leonard is the authority on the subject as he is in a position to confirm James (Stivers et al., 2011).

Hence, it can be seen that even a small excerpt contains many conversational actions. The most prominent action featured was Simone acting as a clarifier. This action was effective as it elucidated important aspects about Leonard’s research and expanded on the topic of effectiveness of the new treatment.
