## Supplementary Table 1 for "Interactional roles in research meetings: a combined conversation analytic and quantitative network approach"

**Supplementary Table 1. Interactional roles and corresponding actions**

| Interactional role | Characteristics | Actions | Example excerpts |
|---|---|---|---|
| **Chair** | ➢ Determines which topics are discussed (both initiating the discussion of some topics and cutting off the discussion of others).<br>➢ Creates space for all group members to express their ideas<br>➢ Encourages group members<br>➢ Keeps the group oriented to the bigger picture | • Monitoring the agenda (both regarding topic and time-concerns) | *And, uh, Tom received a grant, namely the [name grant]. Can we also brainstorm quickly how we're doing on that?* |
| | | • Turning allocation to different group members | *Okay, shall we start with Simone?* |
| | | • Delegating tasks | *[We have] forty minutes. So uh, what about everybody keeps to ten minutes? Is that doable? And can somebody time it?* |
| | | • Orienting to long term goals | *In the background there's always the research question; what exactly is translation? Right, so the- which is something we have been looking into together for years.* |
| | | • Encouraging group members | *Nice going!* |
| **Clarifying** | ➢ Orients to the knowledge of the other<br>➢ Allows others to explain themselves more succinctly or expand on their understanding of the topic<br>➢ Achieves more clarity for the whole group | • Asking questions to clarify or expand on topic | *Where is that receptor then mostly found?* |
| | | • Summarizing topics | *So, the idea is, then, that MS patients have different kinetics than healthy [people], and that is why the axon is more vulnerable to disturbances?* |

| | | | |
|---|---|---|---|
| **Skeptical** | ➢ Asks questions based on their own understanding of the topic<br>➢ Challenges speaker to justify aspects of their research<br>➢ Encourages critical thinking<br>➢ Achieves higher quality research and more clarity concerning specific topics | • Asking challenging questions to ensure higher quality of research | *But can you explain this? Why do you draw this conclusion, then?* |
| **Expert** | ➢ Supports or corrects presenter<br>➢ Is able to close off discussions with their expert knowledge<br>➢ Achieves more clarity for the group | • Intervening when there is confusion | *No, this is actually a linear mixed model, because the GEs are better suited if you also have groups.* |
| **Connecting** | ➢ Facilitates exchange of information<br>➢ Provides a novel point of view, or a novel way of understanding the topic<br>➢ Links different concepts together to form new ideas | • Showcasing the link between different topics<br><br>• Coming up with new ways to conceptualize phenomena | *How does that work compare to what [name] just published?*<br><br>*Could you compare it to a road network, let's say, where some roads are paved and other roads are, uh, cobblestoned, or something like that?* |
| **Practical** | ➢ Supports the presenter by offering concrete advise, help or solutions<br>➢ Showcases group member's willingness to offer help | • Proposing concrete solutions<br><br>• Offering concrete help | *My proposal would be to, if you use the clinical method, or the local method, to determine the outliers, and then correspond that to the worst performance of the rest of the people.*<br><br>*I think she would be super happy to collaborate [with you]. […] I could send an email!* |

*Note.* Provided excerpts are simplified versions of the corresponding occurrences in the data, for the benefit of readability. All examples are underpinned by a detailed conversation analysis, as shown in Supplementary Section 1.
