## Supplementary Table 2 for "Interactional roles in research meetings: a combined conversation analytic and quantitative network approach"

**Supplementary Table 2. Associations of interactional roles to global network measures.**

| | **chair** | | **skeptical** | | **expert** | | **clarifying** | | **connecting** | | **practical** | |
|---|---|---|---|---|---|---|---|---|---|---|---|---|
| | ***r*** | ***p*** | ***r*** | ***p*** | ***r*** | ***p*** | ***r*** | ***p*** | ***r*** | ***p*** | ***r*** | ***p*** |
| **global efficiency** | **-0.683** | **0.001** | 0.073 | 0.897 | **-0.501** | **0.046** | -0.423 | 0.129 | -0.177 | 0.939 | **-0.575** | **0.033** |
| **modularity** | **0.814** | **0.005** | -0.008 | 0.999 | **0.717** | **0.020** | 0.531 | 0.084 | -0.225 | 0.717 | **0.717** | **0.013** |

*Note.* Results were acquired using Pearson's correlation. Significant correlations are presented in bold format. Presented p-values are not corrected for multiple testing.
