## Supplementary Table 3 for "Interactional roles in research meetings: a combined conversation analytic and quantitative network approach"

**Supplementary Table 3. Associations of interactional roles to nodal network measures and meeting evaluations.**

| | | chair | | skeptical | | expert | | clarifying | | connecting | | practical | |
|---|---|---|---|---|---|---|---|---|---|---|---|---|---|
| | | ***β*** | ***p*** | ***β*** | ***p*** | ***β*** | ***p*** | ***β*** | ***p*** | ***β*** | ***p*** | ***β*** | ***p*** |
| ***network measures*** | **local efficiency** | 1.066 | 0.116 | **0,805** | **0.048** | 0.259 | 0.671 | -0.200 | 0.418 | **1.288** | **0.012** | 0.196 | 0.511 |
| | **nodal strength** | **1.211** | **0.012** | 0,364 | 0.212 | 0.156 | 0.702 | -0.283 | 0.099 | **0.784** | **0.025** | 0.015 | 0.941 |
| | **participation coefficient** | 0.499 | 0.446 | -0,242 | 0.596 | 0.939 | 0.186 | 0.278 | 0.293 | 0.371 | 0.543 | -0.435 | 0.224 |
| | **within-module degree** | **1.525** | **0.006** | -0,021 | 0.953 | -0.081 | 0.874 | -0.079 | 0.708 | 0.745 | 0.092 | 0.148 | 0.553 |
| | **betweenness centrality** | **3.328** | **< .001** | 0,052 | 0.886 | 0.818 | 0.109 | -0.307 | 0.152 | 0.252 | 0.575 | -0.319 | 0.206 |
| ***meeting evaluations*** | **1. effectiveness** | 0.238 | 0.354 | -0,196 | 0.355 | 0.063 | 0.853 | -0.069 | 0.483 | -0.221 | 0.292 | -0.076 | 0.506 |
| | **2. worth the investment** | 0.200 | 0.354 | -0,195 | 0.329 | 0.041 | 0.899 | -0.108 | 0.225 | -0.210 | 0.295 | -0.099 | 0.366 |
| | **3. overall experience** | **0.551** | **0.036** | 0,051 | 0.816 | 0.065 | 0.850 | -0.068 | 0.474 | -0.091 | 0.670 | -0.148 | 0.201 |
| | **4. motivating** | 0.040 | 0.853 | -0,050 | 0.799 | -0.073 | 0.810 | -0.047 | 0.581 | -0.211 | 0.259 | 0.046 | 0.658 |
| | **5. educational value** | 0.226 | 0.379 | -0,136 | 0.576 | -0.089 | 0.819 | 0.012 | 0.919 | -0.174 | 0.473 | -0.040 | 0.770 |
| | **6. translational** | 0.262 | 0.281 | -0,067 | 0.765 | 0.438 | 0.200 | 0.059 | 0.545 | -0.226 | 0.295 | -0.046 | 0.697 |
| | **7. feeling of connection** | 0.115 | 0.652 | 0,080 | 0.741 | 0.471 | 0.209 | 0.072 | 0.494 | 0.037 | 0.877 | **-0.305** | **0.014** |
| | **8. able to understand** | 0.077 | 0.775 | 0,124 | 0.642 | -0.268 | 0.529 | -0.019 | 0.866 | 0.193 | 0.467 | -0.209 | 0.142 |

*Note.* Results were acquired using a linear mixed model, accounting for nested data. Significant correlations are presented in bold format. Presented p-values are not corrected for multiple testing. Meeting evaluation items are sequentially ordered and refer to the questionnaire items as listed in Supplementary Section 1.
